# The architecture of metabolic networks constrains the evolution of microbial resource hierarchies

**DOI:** 10.1101/2023.04.21.537789

**Authors:** Sotaro Takano, Jean CC Vila, Ryo Miyazaki, Alvaro Sanchez, Djordje Bajic

## Abstract

Microbial strategies for resource use are an essential determinant of their fitness in complex habitats. When facing environments with multiple nutrients, microbes often use them sequentially according to a preference hierarchy, resulting in well-known patterns of diauxic growth. In theory, the evolutionary diversification of metabolic hierarchies could represent a mechanism supporting coexistence and biodiversity by enabling temporal segregation of niches. Despite this ecologically critical role, the extent to which substrate preference hierarchies can evolve and diversify remains largely unexplored. Here we used genome-scale metabolic modeling to systematically explore the evolution of metabolic hierarchies across a vast space of metabolic network genotypes. We find that only a limited number of metabolic hierarchies can readily evolve, corresponding to the most commonly observed hierarchies in genome-derived models. We further show how the evolution of novel hierarchies is constrained by the architecture of central metabolism, which determines both the propensity to change ranks between pairs of substrates and the effect of specific reactions on hierarchy evolution. Our analysis sheds light on the genetic and mechanistic determinants of microbial metabolic hierarchies, opening new research avenues to understand their evolution, evolvability and ecology.

## Introduction

When presented with an environment with multiple nutrients, many microbes tend to use them one at a time in a preferred order. This phenomenon of hierarchical substrate use was famously characterized by Monod in the decade of 1940 (Monod 1942) when he coined the term “diauxie” to describe the experimentally observed double growth curve pattern. Despite the foundational role of Monod’s work in molecular biology (Jacob and Monod 1961; Belliveau et al. 2018), we still know surprisingly little about the evolution and diversity of resource hierarchies across bacterial species (Perrin et al. 2020). For instance, are some hierarchies easier to evolve than others? And what determines how easy it is to evolve a preference for a given substrate?

The questions about the ecology and evolution of metabolic hierarchies have received renewed attention in recent years (Bajic and Sanchez 2020; Okano et al. 2021), as part of the ongoing effort to understand the drivers of microbial community assembly and coexistence (Chang et al. 2022; Estrela et al. 2022; Gralka et al. 2022). Recent theoretical work (Posfai et al. 2017; Goyal et al. 2018; Pacciani-Mori et al. 2020; Wang et al. 2021; Bloxham et al. 2023) and experiments with model communities (Pacciani-Mori et al. 2020; Bloxham et al. 2022) have shown that differences in metabolic hierarchies can impact ecology, e.g. by allowing species to segregate their metabolic niches and avoid competition. Although these studies carry the implicit assumption that metabolic preferences will readily diversify provided the ecological opportunity (e.g. in an environment with multiple nutrients), it is unclear in which cases this assumption will hold true. Empirical evidence remains scarce and ambiguous. For example, the deep conservation of some preferences, e.g. the almost universal preference for glucose in fermentative microbes (Görke and Stülke 2008), would suggest that metabolic hierarchies are hard to rewire. If this is the case, we would expect metabolic hierarchies to be deeply conserved in the phylogenetic tree and act as a mechanism of coexistence only between distantly related species. However, other studies report divergent resource preferences in closely related species (Tuncil et al. 2017), implying that metabolic hierarchies can quickly diversify. In this scenario, we might expect resource hierarchies to promote the coexistence of closely related strains, and possibly also lead to eco-evolutionary feedbacks (Bajić et al. 2018; Pacciani-Mori et al. 2020). Thus, assessing when and how microbial metabolic hierarchies can diversify is critical to understand their role in structuring coexistence within microbial communities.

A central determinant of the evolution of biological systems is the underlying genotype-phenotype (G-P) map (Fontana and Schuster 1998; Stadler et al. 2001). The architecture of this map determines the amount of genetic variation that can be accessed via mutations, which ultimately fuels evolution. In the case of metabolic traits, the structure of the metabolic network is a central determinant of the G-P map, influencing the evolution of individual enzymes (Papp et al. 2004; Vitkup et al. 2006; Notebaart et al. 2014; Aguilar-Rodríguez and Wagner 2018), metabolic innovation (Barve and Wagner 2013) or eco-evolutionary interactions (Bajić et al. 2018). Importantly, recent work has demonstrated that the structure of the metabolic network is also a key determinant of the strategy microbes adopt in mixed substrate environments, e.g. their choice to use them sequentially versus simultaneously (Wang et al. 2019). Because preferential substrate use represents an optimal metabolic strategy (Salvy and Hatzimanikatis 2021), the presence of one pathway or another will fundamentally determine which substrates an organism prefers, as different pathways have a different balance of benefits and costs when processing a substrate (Noor et al. 2016; Waschina et al. 2016; Wortel et al. 2018). However, how precisely the structure of the metabolic networks determines and constrains the evolution of metabolic hierarchies remains unexplored.

Here, we asked what determines the evolutionary flexibility of microbial metabolic hierarchies using genome-scale metabolic modeling. Metabolic modeling techniques such as flux balance analysis (FBA) enable accurate predictions of metabolic phenotypes from genotypes (Orth et al. 2011; Bordbar et al. 2014; O’Brien et al. 2015) and are widely used as a workhorse for the comprehensive exploration of metabolic genotype-phenotype maps (Segrè et al. 2005; Barve and Wagner 2013; Notebaart et al. 2014; Szappanos et al. 2016; Goldford et al. 2017). Using FBA, we first show that a handful of “typical” resource hierarchies appear much more commonly across genotype space than other configurations. Hierarchies are easier to rewire for substrates that are more metabolically different. However, their evolutionary flexibility strongly depends on the presence or absence of a small number of central metabolic reactions that determine the behavior of the metabolic network across substrates. Our study provides the first systematic analysis of how metabolic networks determine the evolution of metabolic hierarchies, and we end by proposing new testable hypotheses, null expectations and potential directions for future studies.

## Results

### 1 How easily can metabolic hierarchies evolve and diversify?

We began by asking whether metabolic hierarchies could readily evolve and diversify through changes in the set of reactions encoded in a bacterial genome. To address this question, we obtained 9974 random genotypes using random walks through genotype space (Methods). Briefly, to obtain each genotype, we started from the *Escherichia coli* metabolic model *iJO1366* (Orth et al. 2011), and performed 10000 random reaction swaps, removing a reaction from the model and adding another one from a universal set of prokaryotic reactions (Fig. 1A, Methods). This method results in a uniform and unbiased sampling of genotype space (Matias Rodrigues and Wagner 2009).

**Fig. 1.**
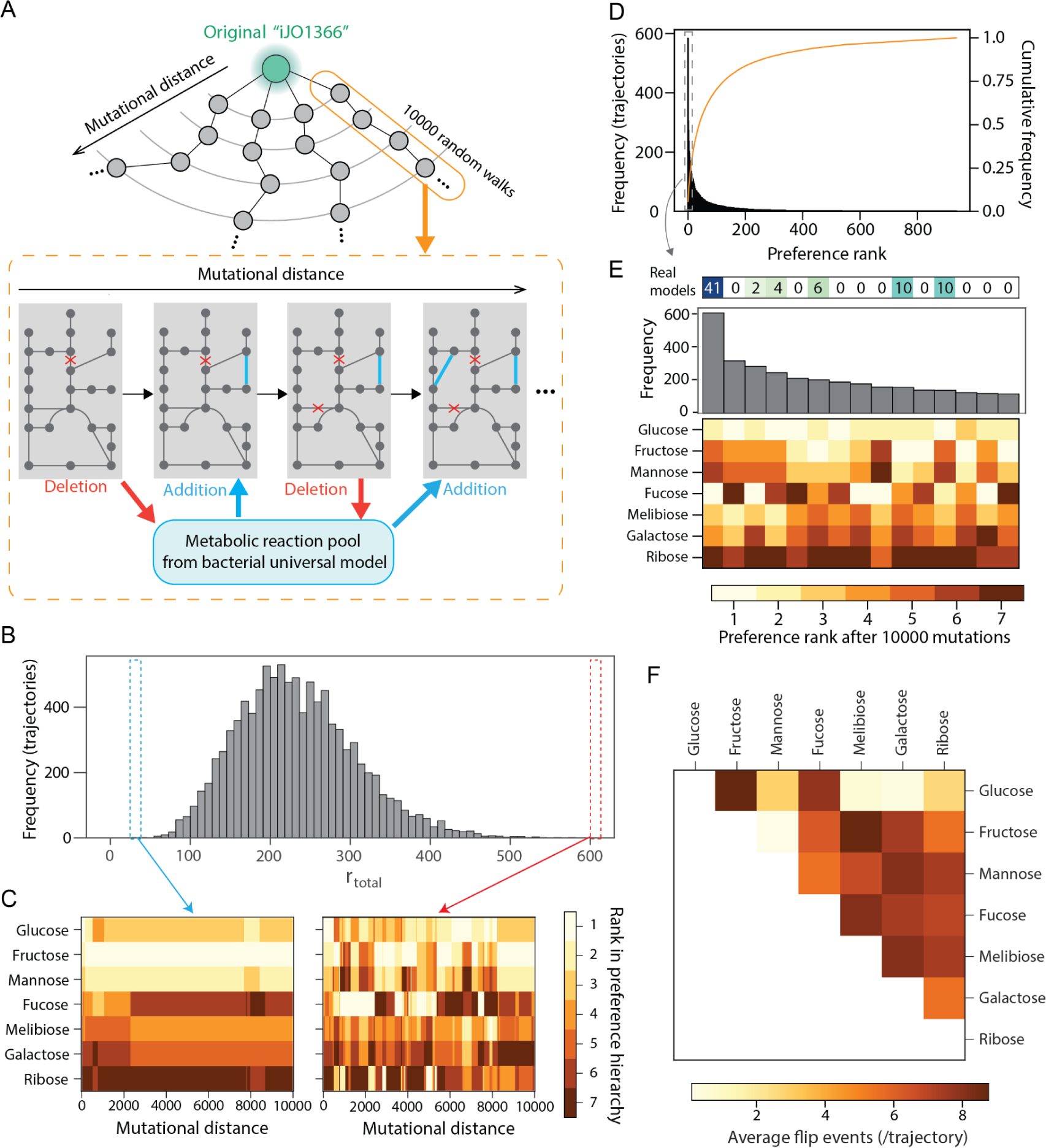
Some sugar hierarchies are easier to evolve than others. (A) A Schematic of random walk trajectories through genotype space. At each step, a reaction from the model is exchanged by a new reaction from a universal bacterial reaction set (“universal model”, Methods). (B) Large variation in metabolic hierarchy rewiring among random walk trajectories. The histogram shows the frequency of total rank flip events (r_total_) during random walks in the 9974 trajectories. (C) Changes in metabolic hierarchy along an example random-walks trajectory. We show two cases, where the rank swaps rarely (left) or frequently (right) occurred. The preference rank of each sugar at each mutational distance is shown as a heatmap. (D) Convergence of the preference rank to small subsets among all possible preference ranks after random walks. The histogram shows the frequency of each preference rank at the end of the random walks in the 9974 evolutionary trajectories. The orange line shows the cumulative distribution of the frequency of preference rank. (E) Zoom of the top 15 most frequent metabolic hierarchies, representing the final point 3069 out of 9974 trajectories (histogram), or about ∼30% of the total. We displayed the number of a real organism’s models whose preference rank matches each rank configuration. (F) Average number of rank flip events between pairs of sugars during random walks in 9974 evolutionary trajectories. Sugars are ordered by their initial preference rank in the *E. coli* model.

We restricted our random walk to the space of genotypes capable of growth on 7 sugars whose growth ranks are well predicted by performing FBA on the *E. coli* model (glucose, fructose, mannose, fucose, melibiose, galactose and ribose, Fig. S1A). Because sequential substrate use is an optimal strategy aimed at maximizing the growth benefit from a substrate (Beg et al. 2007; Kremling et al. 2015; de Groot et al. 2019; Wang et al. 2019; Salvy and Hatzimanikatis 2021), the preference hierarchy usually matches the hierarchy in growth rates (Aidelberg et al. 2014). To confirm this point, we used a global resource allocation constraint (Methods), which is known to reproduce experimentally observed patterns of sequential substrate use (Beg et al. 2007; Salvy and Hatzimanikatis 2021). This confirmed that our substrates are not co-utilized, but sequentially used according to their growth rate hierarchy (Fig. S1B). Thus, for the rest of the paper, we use the growth hierarchy as a proxy for resource use hierarchy in our 7 sugars.

An important driver of the evolution of any trait is the availability of enough phenotypic variation accessible by mutations (Besnard et al. 2020). Even in the presence of selection, it is known that evolution often occurs along genetic lines of maximal variation, or “least resistance” (Schluter 1996). In our trajectories, in absence of selection, rewiring of hierarchies occurred on average fairly often, implying that there is substantial accessible phenotypic variation across the genotype-phenotype map (Fig. 1B and C). However, the outcomes of our trajectories were quite non-uniformly distributed. Out of the 7! = 5040 theoretically possible hierarchy permutations among the 7 sugars, more than 30% of all trajectories (N=9,974) ended in one of ∼15 configurations (Fig. 1D). The outcomes were strongly reminiscent of established empirical observations, e.g. glucose was among the top preferred substrates in many of these configurations. In addition, these typical hierarchies also appeared recurrently in genome-derived models of real organisms (e.g. 73 out of 81 genome-derived models correspond to one of the 7 hierarchies most frequently found in random walk trajectories, Fig. 1E and S3, Methods). This suggests that the metabolic genotype-phenotype map is strongly biased towards a reduced number of resource hierarchy configurations.

A particularly important aspect of hierarchy evolution is the ability to flip ranks between two specific substrates. If two substrates’ ranks can be easily switched, this would allow an easy evolution of metabolic specialization through differential resource preference. When we examined the evolutionary flexibility in the ranks of specific substrate pairs, we found again a very biased distribution in which some sugar pairs swapped their ranks much more often than others in our mutational trajectories (Fig. 1F).

Finally, we also noticed that different trajectories exhibited markedly different degrees of metabolic hierarchy rewiring (mean= 233.2 total hierarchy changes per trajectory, SD = 73.5; Fig. 1B and C, and Fig. S2). This suggests that specific characteristics of the genetic background could modify the amount of accessible variation, and thus the evolutionary flexibility of metabolic hierarchies (Rutherford and Lindquist 1998; Bergman and Siegal 2003; Richardson et al. 2013; Geiler-Samerotte et al. 2019; Poyatos 2020).

Overall, these patterns suggest that the architecture of metabolic networks strongly affects the evolution of metabolic hierarchies by 1) biasing the outcomes of metabolic rewiring towards specific hierarchy configurations, 2) making the rewiring easier for some pairs of substrates than others, and 3) modifying the amount of phenotypic variation accessible to mutations. In the following sections, we study the structural and mechanistic determinants of these patterns.

### 2. Metabolic dissimilarity predicts hierarchy swaps in pairs of substrates

Why are the ranks in some pairs of substrates more easy to change than in others? We reasoned that a primary requirement for two substrates to flip their ranks is the existence of mutations that will affect each of them independently. If two substrates are processed largely by the same pathways, the same mutations should affect their growth similarly (e.g. glucose/melibiose, Fig. 2A) resulting in a lower propensity to flip their ranks (Fig. 2B). In contrast, substrates that are processed by different pathways have more opportunities to evolve independently, and thus swap their hierarchy (e.g. glucose/fucose, Fig. 2C and D). Thus, we hypothesized that the metabolic distance between two substrates will predict their propensity to swap their ranks.

**Fig. 2.**
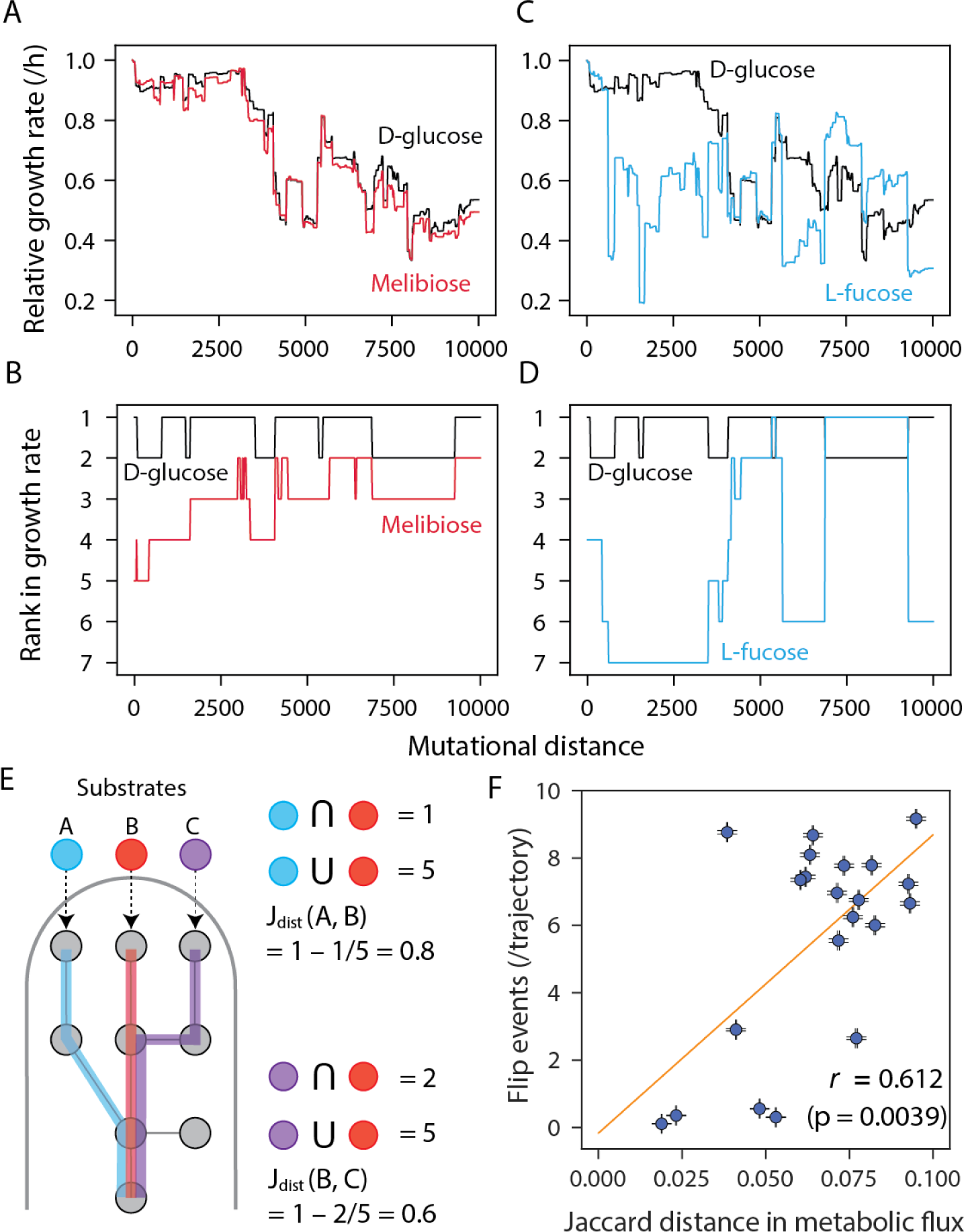
Metabolic dissimilarity predicts hierarchy flips in pairs of substrates. (A) and (C) Representative “random walk” trajectories. Two trajectories are compared for a metabolically similar (D-glucose and Melibiose, A) and dissimilar (D-glucose and L-fucose, C) pairs of sugars. (B) and (D) Resource preference trajectories in the same trajectories shown in A and C. (E) A schematic of calculating Jaccard distance in processing metabolic pathways between two sugars. The number of commonly used reactions for processing both two sugars were divided by the total metabolic reactions used to process each sugar (i.e., Jaccard similarity), and this value was subtracted from 1. In this scheme, we show examples in two pairs of sugars. (F) The dissimilarity in metabolic pathway (Jaccard distance) predicts the rank-flip events in 21 pairs of sugars. An orange line shows linear regressions, error bars indicate +/− SEM, and *r* stands for Pearson’s correlations (N=9974) with Mantel test (10^6^ permutations).

To test this hypothesis we first needed to quantify the metabolic distances between substrate pairs. Because two substrates partially share the pathways and reactions through which they are catabolized, we computed metabolic distance using a weighted Jaccard distance between the sets of reactions used by each metabolite (Fig. 2E, Methods). Although this distance will be genotype-specific (Fig S4A), we observed that the distances between two substrates quickly converged to a typical value across genotypes during random walks (Fig. S4B). Confirming our hypothesis, these “typical” pairwise metabolic distances predicted the propensity of two substrates to rewire their ranks (Fig. 2F). This analysis indicates that the evolutionary flexibility in the hierarchy of two substrates is fundamentally determined by their metabolic dissimilarity.

### 3. The architecture of central metabolism controls the evolutionary flexibility of metabolic hierarchies

As shown in Fig. 1B and C, our random walk trajectories differed markedly in the frequency of shifts in metabolic hierarchies. Although part of this variation is expected because of stochastic sampling of mutations, the presence of specific reactions could also affect the evovability of metabolic hierarchy by either buffering (capacitors) or potentiating (potentiators) the phenotypic effect of other mutations (Rutherford and Lindquist 1998; Bergman and Siegal 2003; Richardson et al. 2013; Geiler-Samerotte et al. 2019; Poyatos 2020). Can we pinpoint the observed differences in the evolutionary flexibility of metabolic hierarchies to the presence or absence of specific metabolic reactions in the genetic background?

To examine this question, we first compared the number of rank shifts (*r_i_*) between trajectories in which a focal reaction was present (*T_R+_*) or absent (*T_R–_*) along the most part of the trajectory (Fig. 3A, Methods). Our analysis revealed 25 reactions which were associated with differences in the evolvability of metabolic rank of at least one sugar (p < 10^−6^, Wilcoxon test with FDR correction, Fig. 3B). To test whether these reactions did indeed alter the phenotypic variation in metabolic hierarchies, we ran two additional sets of 500 random walks, starting from a model either lacking capacitors (“del C”) or potentiators (“del P”, Methods). As expected, these trajectories exhibited an opposite trend in terms of number of rank swap events (Fig. 3C). Thus, the presence of specific metabolic reactions critically determines the evolvability of metabolic hierarchies in our trajectories.

**Fig. 3.**
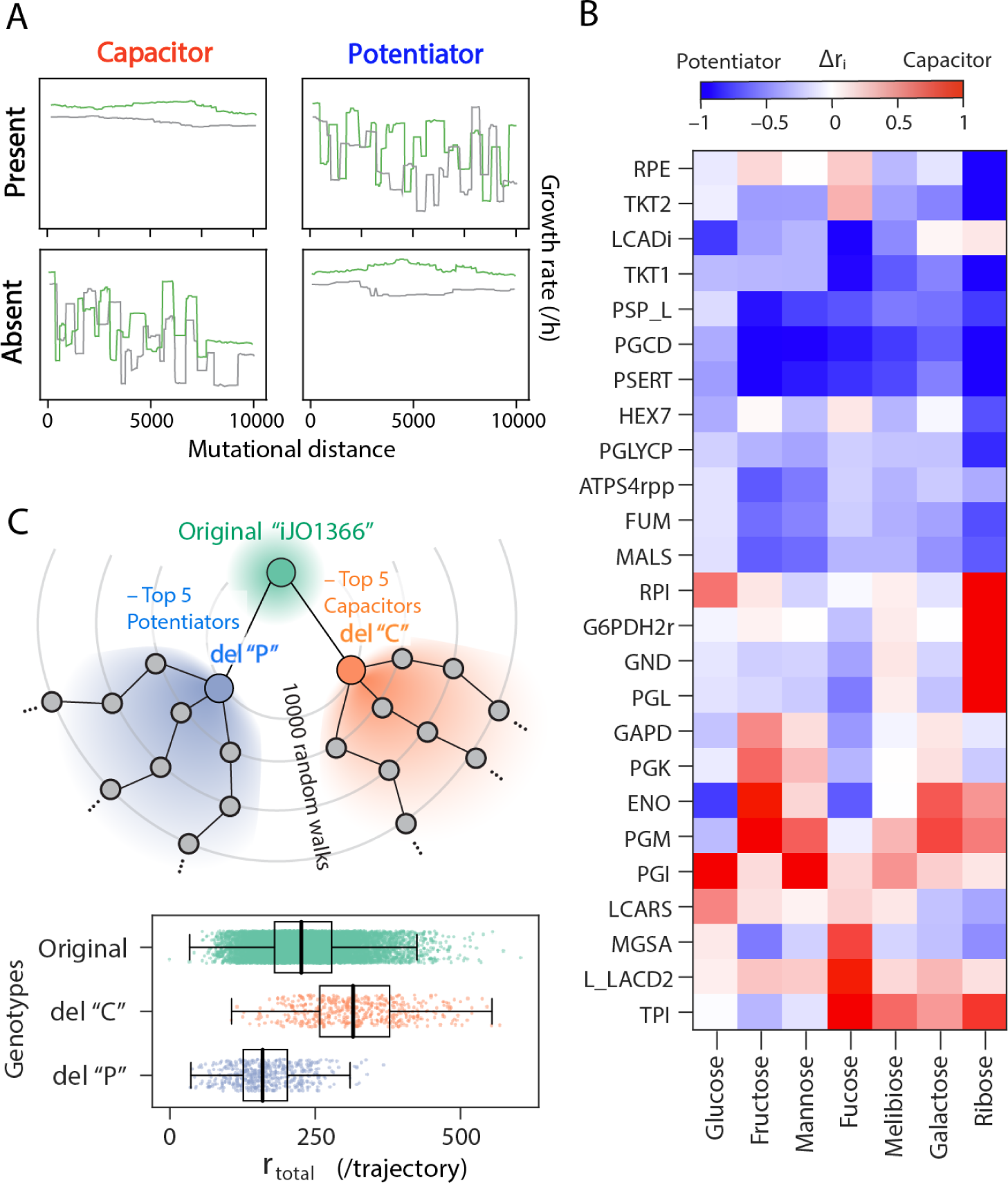
Propensity of rank flips strongly depends on the genetic background. (A) A schematic of the mutational effect of evolutionary capacitors and potentiators on growth-rate trajectories and propensity of rank flips. If the presence of the reaction of interest caused little effect on growth rate and the preference rank (top left) but its deletion caused their frequent changes (bottom left), such reaction was regarded as a “capacitor”. On the other hand, if the growth rate changes and rank flips were prevented by the deletion of the reaction (bottom right) but promoted in its presence, such reaction was regarded as a “potentiator”. Briefly, we screened those evolutionarily important reactions by computing the difference in rank flips (*r_i_*) between its presence (*r_i_* (*T_R+_*)) or absence (*r_i_* (*T_R-_*)) (*Δr_i_*, see Methods). (B) Screened evolutionary capacitors and potentiators in metabolic networks by statistical analysis (p-val < 10^−6^, FDR corrected). *Δr_i_* in each metabolic reaction was shown as a heatmap. The shown metabolic reaction IDs correspond to BiGG ID. (C) Opposite effect on the propensity of rank flips between the deletions of capacitors or potentiators. We deleted 5 potentiators (“del P”) or capacitors (“del C”) from iJO1366 and performed 10000 random walks by starting from each of them. Box and strip plots showed the number of total rank flip events across 7 sugars during random walks (N = 500) in the displayed genetic backgrounds.Bold black lines indicate medians of each data.

### The mechanisms controlling the evolutionary flexibility of metabolic hierarchies

How do specific reactions modify the evolutionary flexibility of metabolic hierarchies? We hypothesized that modifiers act by disproportionately impacting metabolites used by multiple sugars. By doing this, mutations in modifiers can decouple and re-connect sugars with one another, allowing them to evolve more or less independently. This hypothesis is supported by the fact that modifiers generally belong to central metabolism, e.g. glycolysis, tricarboxylic acid (TCA) cycle or pentose phosphate pathway (PPP, Fig. 4A). These reactions tend to carry flux across genotypes and resources (Fig. 4B), and their disruption heavily alters the distribution of fluxes across the rest of the network (Fig. 4C-D).

**Fig. 4.**
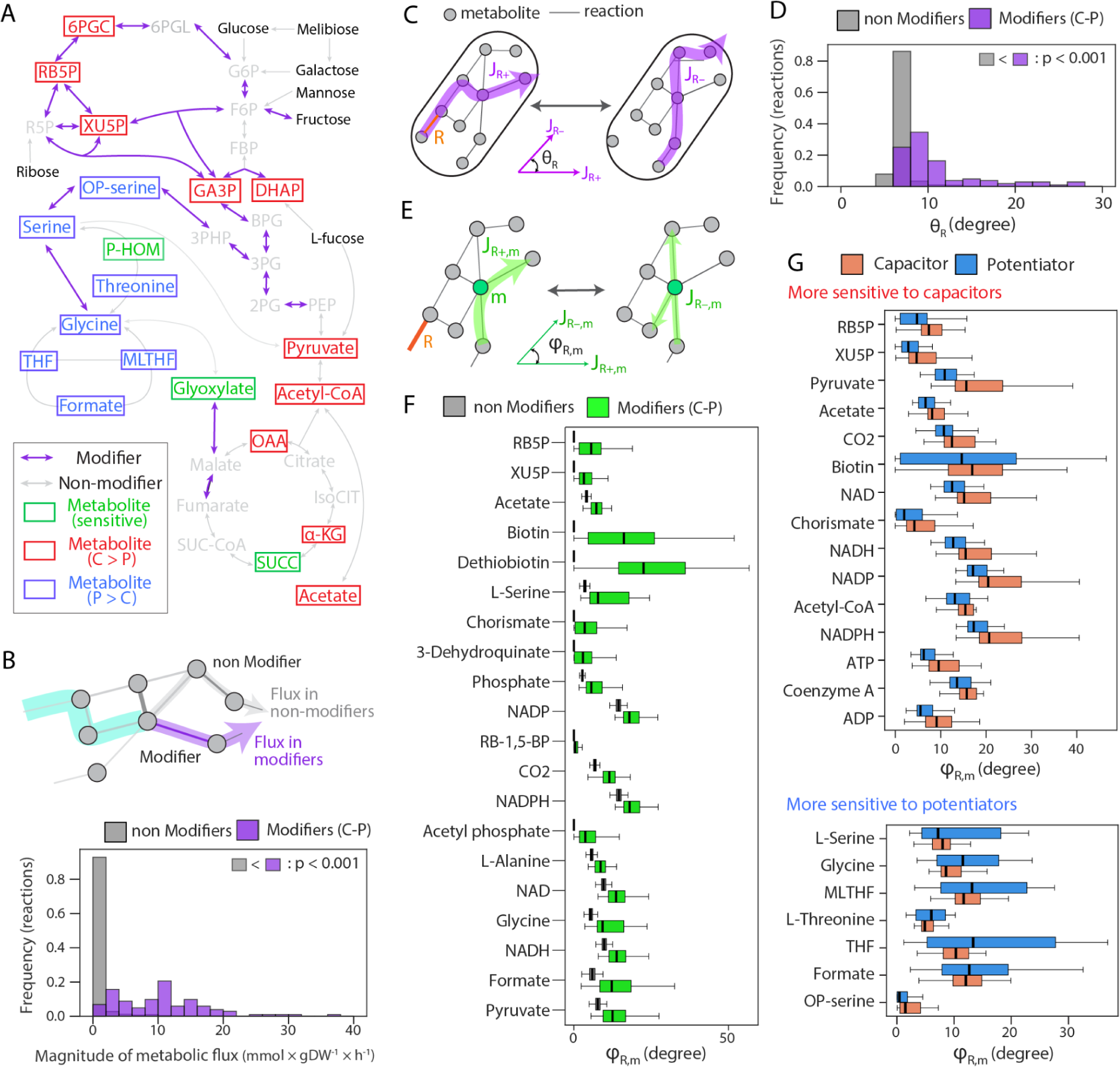
Specific reactions and metabolites govern the effects of mutations on resource hierarchies. (A) The locations of evolutionary modifiers and sensitive metabolites to the modifiers in a metabolic network. Here, we mapped them on the pentose phosphate pathway, glycolysis, serine-glycine pathway, and TCA cycles, where many of those are involved. Metabolites that were more strongly affected by capacitors or potentiators are colored by red and blue, respectively. (B) Comparison of magnitude of flux between modifiers and non-modifiers. We randomly picked up 1000 evolved models and calculated the average flux across 7 sugars in each reaction. Comparison of mean value and statistical significance level were indicated (Wilcoxon ranksum test). (C) A schematic of “flux sensitivity” analysis. For randomly picked up 1000 evolved models, we added or deleted the target reaction *R* depending on whether it exists or not. Then, we calculated the cosine similarity of the total flux distribution between presence (*J_R+_*) or absence (*J_R-_*) of the reaction (*θ_R_*). We calculated *θ_R_*for 7 sugars, and the average of *θ_R_* is used as the proxy for flux sensitivity for the reaction *R.* (D) Comparison of flux sensitivity (*θ_R_*) among non-modifiers (non C-P) or modifiers (C-P). We statistically checked *θ_R_* is significantly larger in modifiers than non-modifiers (Wilcoxon ranksum test, p-value is shown in the panel). (E) A schematic of flux sensitivity analysis for metabolites. For each metabolite *m*, influx and efflux to *m* were analyzed before and after a deletion or addition of a reaction of interest *R*. Then, cosine similarity between *J_R+,m_*and *J_R–,m_* was calculated (*φ_R,m_*). (F) Typical sensitive metabolites to the modifiers. We plotted *φ_R,m_*for top 20 metabolites whose flux were the most significantly affected by mutations of the modifiers. All of the metabolites exhibited significantly higher *φ_R,m_* for modifiers than non-modifiers. Bold black lines indicate medians of each data. (G) Flux sensitivity score (*φ_R,m_*) of metabolites that are differentially affected by capacitors and potentiators are displayed (p < 10^−6^, Wilcoxon ranksum test with FDR correction). Top and bottom panels show the metabolites which were more strongly affected by capacitors and potentiators, respectively. A full name of each metabolite is displayed in Table S1.

To investigate more in detail which specific fluxes are affected by modifiers, we devised a score *φ_R,m_* quantifying the effect of a mutation in reaction *R* on the flux of a metabolite *m* (Fig. 4E). As expected, metabolites associated with central metabolic pathways (glycolysis, pentose phosphate pathway or TCA cycle) were strongly affected (Fig. 4F), especially in the case of capacitors (Fig. 4G). As these pathways are in general the most efficient routes for sugar use, their disruption surfaces other alternatives that might be different for each sugar, thus revealing new mutational targets for the rewiring of metabolic hierarchies. Our analysis also revealed the metabolism of glycine/serine as an important pathway modulating the flexibility of metabolic hierarchies (Fig. 4A and F). Because this pathway can serve as an alternative route from 3-phosphoglycerate to the TCA cycle in our models, its disruption limits available metabolic alternatives. For this reason, metabolites in the glycine/serine pathway appear in general associated with potentiator reactions (Fig. 4G).

Altogether, these results suggest that by shaping the available metabolic alternatives for different substrates and/or their efficiency, the architecture of central metabolism critically determines the evolutionary flexibility of metabolic hierarchies.

## Discussion

The strategies that microorganisms use to metabolize resources have a significant impact on their interactions and coexistence (Bajic and Sanchez 2020; Estrela et al. 2022). Theoretically, having different preferences for the same substrates could enhance biodiversity by allowing temporal niche segregation (Goyal et al. 2018; Bloxham et al. 2023). However, how easily microbial populations evolve alternative metabolic hierarchies remains unclear. In this study, we utilized genome-scale metabolic modeling to investigate how the structure of empirical metabolic genotype-phenotype maps affects the evolution and diversity in the hierarchical use of sugars by microbes. Our findings indicate that the architecture of the metabolic network only permits the evolution of a limited set of hierarchy configurations. Moreover, the evolution of alternative strategies is restricted to substrates that can be processed through substantially different reactions and pathways. Overall, these findings suggest that the diversity of optimal metabolic hierarchies in natural populations may be in general limited.

The evolutionary flexibility of metabolic hierarchies (within the available alternatives) depended on a small set of reactions belonging to central metabolic pathways. One prediction of this result is that sugar hierarchies will show different degrees of conservation across different clades, depending on the architecture of their central carbon metabolism. This suggests a possible explanation to why some phylogenetically distant species show similar metabolic preferences (e.g. the almost universal preference for glucose over other sugars, (Monod 1942; Görke and Stülke 2008)) while at the same time for some species and substrates we find differences between closely related species (Tuncil et al. 2017).

A key assumption of our study is related to optimality in cell behaviour. Firstly, FBA operates under a strong assumption of optimality: phenotypes are predicted by assuming that the kinetic parameters of the metabolic enzymes and their regulation are optimal in a particular environment. While this is generally accepted as a valid approximation (Dykhuizen et al. 1987; Elena and Lenski 2003; Dekel and Alon 2005; Schuetz et al. 2007; Schuetz et al. 2012), it might not be accurate across all conditions (Towbin et al. 2017). Additionally, we assumed that metabolic hierarchies mirror the hierarchy of growth rates. This is reasonable given previous empirical studies (Aidelberg et al. 2014), and more generally, fits the established view that sequential substrate use represents an optimal “economic” strategy that maximizes the benefit obtained from the investment of costly cellular resources in processing a substrate (Beg et al. 2007; Kremling et al. 2015; de Groot et al. 2019; Wang et al. 2019; Salvy and Hatzimanikatis 2021). However, there are possible exceptions to this rule (Okano et al. 2021), e.g. if cells have evolved mechanisms to “prepare” for environmental uncertainty at the cost of optimality in certain environments (Schmidt et al. 2016; Balakrishnan et al. 2021). Deviations from optimality might be especially strong in organisms in which non-metabolic functions (e.g. motility, biofilm formation, persistence) constitute major components of fitness.

An important caveat of our method is the inability to consider the effect of regulatory mutations. For example, *E. coli* implements the preference of some substrates over others through repression of their respective operons at different cAMP concentration thresholds (Okano et al. 2020). Mutations in this regulatory system, e.g. promoter mutations changing the binding strength of the repressor, could therefore represent targets in the evolution of metabolic hierarchies. However, these repression thresholds typically evolve to implement a hierarchy matching the growth rates supported by the substrate. In other words, regulation does not define which metabolic strategy is optimal, but evolves as a means to implementing it. We might therefore expect that regulatory mutations driving the hierarchy away from growth optimality to be typically purged by selection (given that regulatory mechanisms evolve fast compared to their regulation targets (Lozada-Chávez et al. 2006; Price et al. 2008; Aguilar-Rodríguez et al. 2017)). However, exceptions to this rule may emerge under certain ecological contexts. For-example regulatory re-wiring to prefer sub–optimal resources may evolve when resources are supplied sequentially in a non-optimal order or, when an ecological competitor is able to monopolize the most optimal resource.

In summary, our study describes with mechanistic detail how the metabolic genotype-phenotype map influences and constrains the evolution of microbial metabolic hierarchies. This mechanistic perspective has proven to be essential in advancing the field of evolutionary biology, as well as other disciplines (Wagner et al. 2000; de Visser et al. 2003). Future research will be required to explore how the patterns and mechanisms outlined in our study contribute to the phylogenetic and ecological distribution of microbial metabolic hierarchies, as well as their implications for natural and synthetic communities.

## Materials and Methods

### Reconstruction of the genome-scale metabolic model with constrained allocation

We used *E. coli* genome-scale metabolic model *iJO1366* as a reference (Orth et al. 2011). For random walks of metabolic models through genotype space, we constructed bacterial “universal” models as previously described. Briefly, we assembled metabolic reactions in prokaryotic metabolic models posted on the BiGG database, which consists of potential novel reactions in addition to the originally existing reactions in the *E. coli* model. We modified the directionality of metabolic reactions and removed erroneous energy-generating cycles as previously described. This generates the prokaryotic “universal” models, consisting of 5584 reactions and 3476 metabolites.

Because the cell has limited internal resources, inefficient but “cheap” pathways result in a higher growth rate than more efficient but expensive ones, by allowing for allocation of larger fractions of the proteome to uptake (Basan et al. 2015). We partially account for this using a global resource allocation constraint (CAFBA (Mori et al. 2016)) and saturating sugar concentrations. To this end, we implemented the CAFBA constraint as in the original paper. Briefly, partitioning of cellular resources (i.e. proteome) associated with ribosome, biosynthetic enzymes, carbon transport, and basic biological processes (i.e. housekeeping reactions) was introduced as *φ_R_, φ_E_, φ_C_,* and *φ_Q_*, respectively. By the empirical studies, the sum of those proteome fractions should be 1 (i.e. *φ_R_* + *φ_E_ + φ_C_* + *φ_Q_* = 1), and each fraction except for *φ_Q_* can be altered by environmental conditions and physiological states. Previous studies phenomenologically demonstrated dependencies of each fraction on environmental factors: ribosome sector *φ_R_* is proportionally changed by the growth rate *λ* (i.e. *Δφ_R_* = *w_R_λ*); biosynthetic enzymes sector *φ_E_* changes proportional to the metabolic flux *v*_i_ such that *Δφ_E_* = ∑_i_*w_i_|v*_i_|; carbon transport sector *φ_C_* alters proportional to the carbon intake flux *v*_c_ (i.e. *Δφ_c_*= *w_c_v_c_*) . CAFBA maximizes objective function while optimally partitioning those flexible proteome fraction, *φ_max_* = *Δφ_R_* + *Δφ_E_* + *Δφ_C_*. By incorporating those schemes to FBA, we computed flux distribution to maximize growth rate *λ* with optimally allocating cellular resources to those three partitions. Then the optimization problem of CAFBA is formulated as follows.

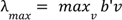

subject to *Sv* = 0

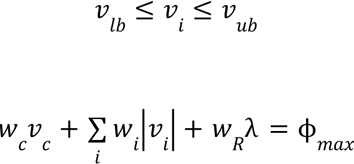

where λ_max_ denotes the maximum growth rate, and *v* is a metabolic flux vector with the lower and upper bounds (i.e. *v_lb_* and *v_ub_*). *b* is the vector of objective function. *S* denotes the stoichiometric matrix of metabolic networks. *w_R_* and *w_c_*is a coefficient for *φ_R_*and *φ_C_*, respectively. ∑_i_*w_i_|v*_i_| is the sum in metabolic flux catalyzed by enzymes except for transport and exchange reactions.

### Metabolic simulation of growth

All FBA simulations were performed using the COBRApy package (Ebrahim et al. 2013). To simulate the growth of bacteria where the limiting factor is only the carbon source, we ran all the simulations under the condition that inorganic ions and gases (ca2_e, cl_e, cobalt2_e, cu2_e, fe2_e, fe3_e, h2O_e, h_e, k_e, mg2_e, mn2_e, mobd_e, na1_e, nh4_e, ni2_e, pi_e, so4_e, zn2_e, o2_e) were present in excess (i.e. the lower bound for exchange reactions for those metabolites was set to –1000 mmol×gDW^−1^×h^−1^). The lower bound of the exchange reaction for each sugar is set so that the influx in C atoms is −120 mmol×gDW^−1^×h^−1^ (i.e. if the given sugar is a hexose such as glucose, there is six carbon atoms per molecule, so the lower limit is set to −20 mmol×gDW^−1^×h^−1^). In all simulations, the objective function is the biomass function of *iJO1366* (i.e. BIOMASS_Ec_iJO1366_core_53p95M). Optimization problems were solved with Gurobi or CPLEX optimizer.

### Simulation of the repressive effect on sugar influx by metabolic hierarchy

The repressive effect on influx of sugar (i.e. flux in exchange reaction) by the presence of other sugar was simulated by parsimonious FBA (pFBA, Fig. S1). The influx of sugar *i,* where no other sugars present is first computed, which is denoted as *J_i_*. Then the influx in sugar *i* in the presence of sugar *j* is simulated. The repressive effect of sugar *j* on the influx of sugar *i,* which is denoted as *R_i,j_* is formulated as follows:

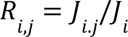

*J_i_* should always be larger than *J_i,j_*, because the influx of sugar generally should be maximal when that sugar is used as a sole carbon source for the growth. Then, *R_i,j_* < 1. If *R_i,j_* ≈ 1, the influx in sugar *i* was barely affected by sugar *j,* indicating that sugar *i* is ranked higher than sugar *j* in metabolic hierarchy. On the other hand, if *R_i,j_* ≈ 0, the influx of sugar *i* is strongly repressed by the presence of sugar *j*, indicating that sugar *i* is ranked lower than sugar *j.* The lower bound of the exchange reaction for sugars is set so that the influx in Catoms is −120 mmol×gDW^−1^×h^−1^.

### Random walks in genotype map

To explore the evolvability of metabolic hierarchy in sugars, we performed random walk by deleting and adding metabolic reactions one by one starting from the *iJO1366* model. Since we focused on the effect of mutations in the intracellular metabolic network, transport, exchange, and sink reactions were not deleted or added and remained as original. The prokaryotic metabolic reaction pool, candidates for newly appended reactions, was prepared by selecting reactions in a universal model which does not exist in the original *iJO1366* model. In each step of random walks, we randomly deleted an existing metabolic reaction in a model and then randomly appended the novel reaction from the prokaryotic reaction pool as long as the model can utilize all 7 sugars as a growth-substrate (i.e. grow on each sugar as a sole carbon source). Once the reaction-swap event was accomplished, the deleted reaction from a model was added to the members of the prokaryotic reaction pool and regarded as a novel reaction. On the other hand, the appended reaction to the model was removed from the prokaryotic reaction pool (Fig. 1A). This random swap was performed 5000 times, resulting in 10000 additions and deletions of reactions. During random walks, the coexistence of following pairs of reactions was avoided: SHSL2 and SHSL2r, DHORD_NAD and DHORDi, ENO and HADPCOADH, LEUTA and LLEUDr, P5CRx and PRO1y, because it leads to CO_2_ or H_2_ limitation (Bajić et al. 2018).

### Estimation of the rank in metabolic hierarchy by growth-rate

Since the ranks of 7 sugars in the metabolic hierarchy correspond to the ranks in growth rate when each sugar is used as a sole carbon source (for details, see Fig. S1 and Supporting Text), we computed the growth rate on every 7 sugars during random walks and used this metric for estimating the rank in the hierarchy. We compared the growth rate on each sugar and regarded that sugar *i* is ranked higher than sugar *j* if the log10 ratio of the growth rate on sugar *i* (denoted as *g*_i_) and *j* (denoted as *g*_j_) is more than 10^−5^ as follows:

For sugar *i,j,* < *j* in the metabolic hierarchy,

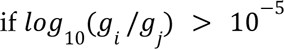

### Computation of preference ranks in genome-derived models

Using the same criteria as above, we computed the rank in the metabolic hierarchy in the prokaryotic metabolic models constructed by CarveMe pipeline. We started from a sample of 5587 metabolic models available at https://github.com/cdanielmachado/embl_gems (Machado et al. 2018). To obtain a meaningful statistical sample, we searched for the set of sugars able to individually support the growth of a maximum number of models. This resulted in 5 sugars (glucose, fructose, mannose, galactose, ribose) that were able to support the growth of 81 models in total (glucose, fructose, mannose, galactose, ribose). These models are taxonomically classified as 3 phyla (*Firmicutes, Proteobacteria,* and *Actinobacteria*). In Fig. S3B, we performed a random permutation test to check the statistical significance levels of Pearson’s correlation in the frequency of rank configurations between random-walks and real organism’s metabolic reconstructions. We shuffled the frequency in random walks of rank configurations (left in panel A and y-axis of panel B), and computed the Pearson’s correlation using those shuffled data for 100000 times. P-value is the probability that the correlations between shuffled data were larger than that observed in the original data.

### Computation of dissimilarity between pairs of sugars in processing metabolic pathways

We estimated dissimilarity in used metabolic pathways between two sugars by calculating Jaccard distance in the set of active reactions where the sugar of interest is used for growth as a sole carbon source. We first simulated the flux distribution in intracellular metabolic reactions (i.e. transport reactions were not included) when the model grows on sugar *i* by pFBA. Then the set of active reactions ***R***, whose flux is more than 10^−6^ mmol×gDW^−1^×h^−1^, was picked out. The Jaccard distance between sugar *i* and *j* in the processing pathways (*J_dist_*(*i,j*)) was formulated as follows:

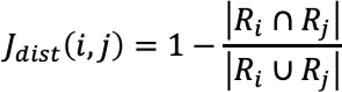

### Screening of capacitors and potentiators

To screen the reactions whose presence or absence greatly affect the evolvability of the hierarchy during random walks, we selected two types of random walks trajectories: *T_R+_,* where the reaction of interest *R* over 70 % of genotypes among first 7000 mutations; *T_R–_*, where the reaction of interest was not present over 70 % of genotypes among last 7000 mutations. Then we statistically tested whether *r_i_* (propensity of rank flips) was significantly different between *T_R+_* and *T_R–_* for each reaction *R* by *t-test* using *scipy.stats.ttest_ind* function with p-value correction for multiple tests by holm-sidak method using *statsmodels.stats.multitest.multipletests*. We screened the reaction showing p-value < 10^−6^ after correction. We also calculated the magnitude of changes in *r_i_* between *T_R+_*and *T_R–_* as follows.

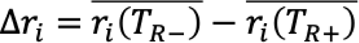

If *Δr_i_* is positive, the absence of reaction *R* increases the evolutionary flexibility of metabolic hierarchies of the hierarchy by mutations, which means that the reaction *R,* as it were, masks the effect of mutations during its presence. Then such a reaction is defined as a “capacitor” for rewiring the metabolic hierarchy. On the other hand, if *Δr_i_* is negative, the presence of reaction *R* increases the propensity of rewiring the hierarchy, and thus is defined as a “potentiator”. We screened such reactions for two metrics and for each 7 sugars.

### Random walks using the models lacking capacitors or potentiators

To confirm the effect of modifiers on the evolutionary flexibility of the metabolic hierarchy, we removed 5 representative capacitors or 5 potentiators from the original *iJO1366* before performing additional random walks. We first selected reactions which solely work as either capacitor or potentiator (i.e., not dual ways like “ENO”, which work as a potentiator for glucose but also work as a capacitor for fructose). In each case of potentiator or capacitor, the reactions were sorted by the number of significantly affected substrates. Then the reactions were deleted from the *iJO1366* continuously from the top of the list up to 5 as long as the model can grow on all 7 sugars (i.e., If the reaction is essential for the growth on either of 7 sugars, that deletion is canceled). This gives us the models lacking 5 capacitors, “PGM”, “PGI”, ”TPI”, ”L_LACD2”, ”LCARS”, (“del C” in Fig. 3C) or 5 potentiators, “PSERT”, “PGCD”, “PSP_L”, “FUM”, “MALS” (“del P” in Fig. 3C) and being capable of utilizing all 7 sugars. Then, we performed 500 independent random walks in each case.

### Flux sensitivity analysis

The impact of mutation to reaction *i* (i.e. deletion or addition of the reaction) on the intracellular metabolic flux is estimated by calculating cosine similarity in the flux distribution between before and after the mutation. For randomly mutated models (i.e. models after 10000 random walks in genotype space), we added or deleted the reaction of interest (denoted as *R*) if that reaction is absent or present, respectively. We first computed the intracellular flux distribution by pFBA before or after the mutation to *R*, which is denoted as *J_R+_*or *J_R-_*, and normalized it by the growth rate (i.e., the magnitude of biomass flux) as follows.

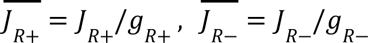

Here, *g_R+_* and *g_R–_* are growth rates in the presence or absence of reaction *R,* respectively. Then we computed the cosine similarity between those normalized flux distribution matrices as follows:

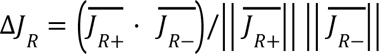

As a metric of the distance, we used *θ_R_* (degree). *ΔJ_R_* is converted to *θ_R_* as follows:

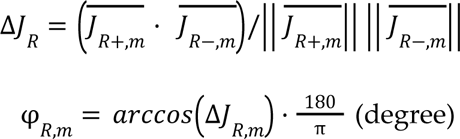

We computed *θ_R_* across 7 sugars and the average value of that is used as the metric for flux sensitivity.

For computing the impact on the metabolic flux in a specific metabolite, we picked out fluxes of reactions where the target metabolite *m* is involved from the normalized intracellular metabolic flux, which are denoted as 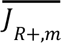 and 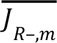, respectively. Then the impact of mutation to reaction *R* on the flux in metabolite *m* is as follows:

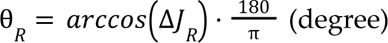

### Experimental quantification of growth hierarchy in *Escherichia coli*

Escherichia coli MG1655 was streaked from glycerol on a TSA plate and grown at 37C for 24h. A single colony was used to inoculate 5mL of TSB medium in a 50mL Falcon tube. After 24h incubation at 37C, this pre-culture was then diluted 1:1000 in M9 minimal medium supplemented with each of 7 carbon sources (glucose, fructose, mannose, fucose, melibiose, galactose and ribose), at a final concentration of 0.07 moles of carbon per liter. *E. coli* growth was monitored in a 384-well plate (100uL/well, 6 replicates each) at 37C by optical density (OD) measurements. The maximal exponential growth rate was computed by first smoothing the log(OD620) with a generalized additive model with an adaptive smoother, using the gam function from the mgcv package in R. This method allows for extraction of estimates of growth rate that are not biased by underlying assumptions when fitting parametric models such as logistic or Gompertz. The maximum of the derivative was taken as the exponential growth rate. The first 1h of growth as well as all of the timepoints in the beginning of the curve that showed an OD<0.01 were excluded to avoid artifacts derived from measurement and fitting noise.

## Acknowledgements

We want to thank members of the Sanchez, Bajic and Miyazaki groups for their helpful discussion.

## Funding

This work was partially funded by Young Investigator Award RGY0077/2016 from the Human Frontier Science Program to A.S and R.M. A.S. was partially supported by the Spanish Ministry of Science and Innovation, under project PID2021-125478NA-100. S.T. was supported by Japan Society for the Promotion of Science postdoctoral fellowship (18J01280).

## Author contributions

D.B. and A.S. conceived the idea and designed the study. S.T. carried out the simulations, formal analysis and figure preparation. J.C.C.V, R.M. provided materials and technical assistance. S.T., J.C.C.V, R.M., A.S, D.B. discussed the results and drafted the paper. S.T. and D.B. wrote the final version of the paper.

## Competing interests

The authors declare no competing interests.

## Data and materials availability

All code for simulations, data analysis and figures can be found in the following github repository https://github.com/sotarotakano/MetabolicHierarchy. Data and code will be provided with corresponding DOI before publication.

## Notes

### Competing Interest Statement

The authors have declared no competing interest.

## References

Aguilar-Rodríguez J, Payne JL, Wagner A. 2017. A thousand empirical adaptive landscapes and their navigability. Nat Ecol Evol 1:45.

Aguilar-Rodríguez J, Wagner A. 2018. Metabolic Determinants of Enzyme Evolution in a Genome-Scale Bacterial Metabolic Network. Genome Biol. Evol. 10:3076–3088.

Aidelberg G, Towbin BD, Rothschild D, Dekel E, Bren A, Alon U. 2014. Hierarchy of non-glucose sugars in Escherichia coli. BMC Syst. Biol. 8:133.

Bajic D, Sanchez A. 2020. The ecology and evolution of microbial metabolic strategies. Curr. Opin. Biotechnol. 62:123–128.

Bajić D, Vila JCC, Blount ZD, Sánchez A. 2018. On the deformability of an empirical fitness landscape by microbial evolution. Proc. Natl. Acad. Sci. U. S. A. 115:11286–11291.

Balakrishnan R, de Silva RT, Hwa T, Cremer J. 2021. Suboptimal resource allocation in changing environments constrains response and growth in bacteria. Mol. Syst. Biol. 17:e10597.

Barve A, Wagner A. 2013. A latent capacity for evolutionary innovation through exaptation in metabolic systems. Nature 500:203–206.

Basan M, Hui S, Okano H, Zhang Z, Shen Y, Williamson JR, Hwa T. 2015. Overflow metabolism in Escherichia coli results from efficient proteome allocation. Nature 528:99–104.

Beg QK, Vazquez A, Ernst J, de Menezes MA, Bar-Joseph Z, Barabási A-L, Oltvai ZN. 2007. Intracellular crowding defines the mode and sequence of substrate uptake by Escherichia coli and constrains its metabolic activity. Proc. Natl. Acad. Sci. U. S. A. 104:12663–12668.

Belliveau NM, Barnes SL, Ireland WT, Jones DL, Sweredoski MJ, Moradian A, Hess S, Kinney JB, Phillips R. 2018. Systematic approach for dissecting the molecular mechanisms of transcriptional regulation in bacteria. Proc. Natl. Acad. Sci. U. S. A. 115:E4796–E4805.

Bergman A, Siegal ML. 2003. Evolutionary capacitance as a general feature of complex gene networks. Nature 424:549–552.

Besnard F, Picao-Osorio J, Dubois C, Félix M-A. 2020. A broad mutational target explains a fast rate of phenotypic evolution. Elife [Internet] 9. Available from: http://dx.doi.org/10.7554/eLife.54928

Bloxham B, Lee H, Gore J. 2022. Diauxic lags explain unexpected coexistence in multi-resource environments. Mol. Syst. Biol. 18:e10630.

Bloxham B, Lee H, Gore J. 2023. Biodiversity is enhanced by sequential resource utilization and environmental fluctuations via emergent temporal niches. bioRxiv [Internet]:2023.02.17.529002. Available from: https://www.biorxiv.org/content/10.1101/2023.02.17.529002v1.abstract

Bordbar A, Monk JM, King ZA, Palsson BO. 2014. Constraint-based models predict metabolic and associated cellular functions. Nat. Rev. Genet. 15:107–120.

Chang C-Y, Bajic D, Vila J, Estrela S, Sanchez A. 2022. Emergent coexistence in multispecies microbial communities. bioRxiv [Internet]:2022.05.20.492860. Available from: https://www.biorxiv.org/content/10.1101/2022.05.20.492860v2.abstract

Dekel E, Alon U. 2005. Optimality and evolutionary tuning of the expression level of a protein. Nature 436:588–592.

Dykhuizen DE, Dean AM, Hartl DL. 1987. Metabolic flux and fitness. Genetics 115:25–31.

Ebrahim A, Lerman JA, Palsson BO, Hyduke DR. 2013. COBRApy: COnstraints-Based Reconstruction and Analysis for Python. BMC Syst. Biol. 7:74.

Elena SF, Lenski RE. 2003. Evolution experiments with microorganisms: the dynamics and genetic bases of adaptation. Nat. Rev. Genet. 4:457–469.

Estrela S, Vila JCC, Lu N, Bajić D, Rebolleda-Gómez M, Chang C-Y, Goldford JE, Sanchez-Gorostiaga A, Sánchez Á. 2022. Functional attractors in microbial community assembly. Cell Syst 13:29–42.e7.

Fontana W, Schuster P. 1998. Shaping space: the possible and the attainable in RNA genotype-phenotype mapping. J. Theor. Biol. 194:491–515.

Geiler-Samerotte K, Sartori FMO, Siegal ML. 2019. Decanalizing thinking on genetic canalization. Semin. Cell Dev. Biol. 88:54–66.

Goldford JE, Hartman H, Smith TF, Segrè D. 2017. Remnants of an Ancient Metabolism without Phosphate. Cell 168:1126–1134.e9.

Görke B, Stülke J. 2008. Carbon catabolite repression in bacteria: many ways to make the most out of nutrients. Nat. Rev. Microbiol. 6:613–624.

Goyal A, Dubinkina V, Maslov S. 2018. Multiple stable states in microbial communities explained by the stable marriage problem. ISME J. 12:2823–2834.

Gralka M, Pollak S, Cordero OX. 2022. Fundamental metabolic strategies of heterotrophic bacteria. bioRxiv [Internet]:2022.08.04.502823. Available from: https://www.biorxiv.org/content/10.1101/2022.08.04.502823v1.abstract

de Groot DH, van Boxtel C, Planqué R, Bruggeman FJ, Teusink B. 2019. The number of active metabolic pathways is bounded by the number of cellular constraints at maximal metabolic rates. PLoS Comput. Biol. 15:e1006858.

Jacob F, Monod J. 1961. Genetic regulatory mechanisms in the synthesis of proteins. J. Mol. Biol. 3:318–356.

Kremling A, Geiselmann J, Ropers D, de Jong H. 2015. Understanding carbon catabolite repression in Escherichia coli using quantitative models. Trends Microbiol. 23:99–109.

Lozada-Chávez I, Janga SC, Collado-Vides J. 2006. Bacterial regulatory networks are extremely flexible in evolution. Nucleic Acids Res. 34:3434–3445.

Machado D, Andrejev S, Tramontano M, Patil KR. 2018. Fast automated reconstruction of genome-scale metabolic models for microbial species and communities. Nucleic Acids Res. 46:7542–7553.

Matias Rodrigues JF, Wagner A. 2009. Evolutionary plasticity and innovations in complex metabolic reaction networks. PLoS Comput. Biol. 5:e1000613.

Monod J. 1942. Recherches sur la croissance des cultures bacteriennes. (Hermann & Cie, editor.). Paris

Mori M, Hwa T, Martin OC, De Martino A, Marinari E. 2016. Constrained Allocation Flux Balance Analysis. PLoS Comput. Biol. 12:e1004913.

Noor E, Flamholz A, Bar-Even A, Davidi D, Milo R, Liebermeister W. 2016. The Protein Cost of Metabolic Fluxes: Prediction from Enzymatic Rate Laws and Cost Minimization. PLoS Comput. Biol. 12:e1005167.

Notebaart RA, Szappanos B, Kintses B, Pál F, Györkei Á, Bogos B, Lázár V, Spohn R, Csörgő B, Wagner A, et al. 2014. Network-level architecture and the evolutionary potential of underground metabolism. Proc. Natl. Acad. Sci. U. S. A. 111:11762–11767.

O’Brien EJ, Monk JM, Palsson BO. 2015. Using Genome-scale Models to Predict Biological Capabilities. Cell 161:971–987.

Okano H, Hermsen R, Hwa T. 2021. Hierarchical and simultaneous utilization of carbon substrates: mechanistic insights, physiological roles, and ecological consequences. Curr. Opin. Microbiol. 63:172–178.

Okano H, Hermsen R, Kochanowski K, Hwa T. 2020. Regulation underlying hierarchical and simultaneous utilization of carbon substrates by flux sensors in Escherichia coli. Nat Microbiol 5:206–215.

Orth JD, Conrad TM, Na J, Lerman JA, Nam H, Feist AM, Palsson BØ. 2011. A comprehensive genome-scale reconstruction of Escherichia coli metabolism--2011. Mol. Syst. Biol. 7:535.

Pacciani-Mori L, Giometto A, Suweis S, Maritan A. 2020. Dynamic metabolic adaptation can promote species coexistence in competitive microbial communities. PLoS Comput. Biol. 16:e1007896.

Papp B, Pál C, Hurst LD. 2004. Metabolic network analysis of the causes and evolution of enzyme dispensability in yeast. Nature 429:661–664.

Perrin E, Ghini V, Giovannini M, Di Patti F, Cardazzo B, Carraro L, Fagorzi C, Turano P, Fani R, Fondi M. 2020. Diauxie and co-utilization of carbon sources can coexist during bacterial growth in nutritionally complex environments. Nat. Commun. 11:3135.

Posfai A, Taillefumier T, Wingreen NS. 2017. Metabolic Trade-Offs Promote Diversity in a Model Ecosystem. Phys. Rev. Lett. 118:028103.

Poyatos JF. 2020. Genetic buffering and potentiation in metabolism. PLoS Comput. Biol. 16:e1008185.

Price MN, Dehal PS, Arkin AP. 2008. Horizontal gene transfer and the evolution of transcriptional regulation in Escherichia coli. Genome Biol. 9:R4.

Richardson JB, Uppendahl LD, Traficante MK, Levy SF, Siegal ML. 2013. Histone variant HTZ1 shows extensive epistasis with, but does not increase robustness to, new mutations. PLoS Genet. 9:e1003733.

Rutherford SL, Lindquist S. 1998. Hsp90 as a capacitor for morphological evolution. Nature 396:336–342.

Salvy P, Hatzimanikatis V. 2021. Emergence of diauxie as an optimal growth strategy under resource allocation constraints in cellular metabolism. Proc. Natl. Acad. Sci. U. S. A. [Internet] 118. Available from: http://dx.doi.org/10.1073/pnas.2013836118

Schluter D. 1996. Adaptive radiation along genetic lines of least resistance. Evolution 50:1766–1774.

Schmidt A, Kochanowski K, Vedelaar S, Ahrné E, Volkmer B, Callipo L, Knoops K, Bauer M, Aebersold R, Heinemann M. 2016. The quantitative and condition-dependent Escherichia coli proteome. Nat. Biotechnol. 34:104–110.

Schuetz R, Kuepfer L, Sauer U. 2007. Systematic evaluation of objective functions for predicting intracellular fluxes in Escherichia coli. Mol. Syst. Biol. 3:119.

Schuetz R, Zamboni N, Zampieri M, Heinemann M, Sauer U. 2012. Multidimensional optimality of microbial metabolism. Science 336:601–604.

Segrè D, Deluna A, Church GM, Kishony R. 2005. Modular epistasis in yeast metabolism. Nat. Genet. 37:77–83.

Stadler BM, Stadler PF, Wagner GP, Fontana W. 2001. The topology of the possible: formal spaces underlying patterns of evolutionary change. J. Theor. Biol. 213:241–274.

Szappanos B, Fritzemeier J, Csörgő B, Lázár V, Lu X, Fekete G, Bálint B, Herczeg R, Nagy I, Notebaart RA, et al. 2016. Adaptive evolution of complex innovations through stepwise metabolic niche expansion. Nat. Commun. 7:11607.

Towbin BD, Korem Y, Bren A, Doron S, Sorek R, Alon U. 2017. Optimality and sub-optimality in a bacterial growth law. Nat. Commun. 8:14123.

Tuncil YE, Xiao Y, Porter NT, Reuhs BL, Martens EC, Hamaker BR. 2017. Reciprocal Prioritization to Dietary Glycans by Gut Bacteria in a Competitive Environment Promotes Stable Coexistence. MBio [Internet] 8. Available from: http://dx.doi.org/10.1128/mBio.01068-17

de Visser JAGM, Hermisson J, Wagner GP, Ancel Meyers L, Bagheri-Chaichian H, Blanchard JL, Chao L, Cheverud JM, Elena SF, Fontana W, et al. 2003. Perspective: Evolution and detection of genetic robustness. Evolution 57:1959–1972.

Vitkup D, Kharchenko P, Wagner A. 2006. Influence of metabolic network structure and function on enzyme evolution. Genome Biol. 7:R39.

Wagner GP, Chiu C-H, Laubichler M. 2000. Developmental Evolution as a Mechanistic Science: The Inference from Developmental Mechanisms to Evolutionary Processes. Integr. Comp. Biol. 40:819–831.

Wang X, Xia K, Yang X, Tang C. 2019. Growth strategy of microbes on mixed carbon sources. Nat. Commun. 10:1279.

Wang Z, Goyal A, Dubinkina V, George AB, Wang T, Fridman Y, Maslov S. 2021. Complementary resource preferences spontaneously emerge in diauxic microbial communities. Nat. Commun. 12:6661.

Waschina S, D’Souza G, Kost C, Kaleta C. 2016. Metabolic network architecture and carbon source determine metabolite production costs. FEBS J. 283:2149–2163.

Wortel MT, Noor E, Ferris M, Bruggeman FJ, Liebermeister W. 2018. Metabolic enzyme cost explains variable trade-offs between microbial growth rate and yield. PLoS Comput. Biol. 14:e1006010.

