## supplement_figs_and_text for "The architecture of metabolic networks constrains the evolution of microbial resource hierarchies"

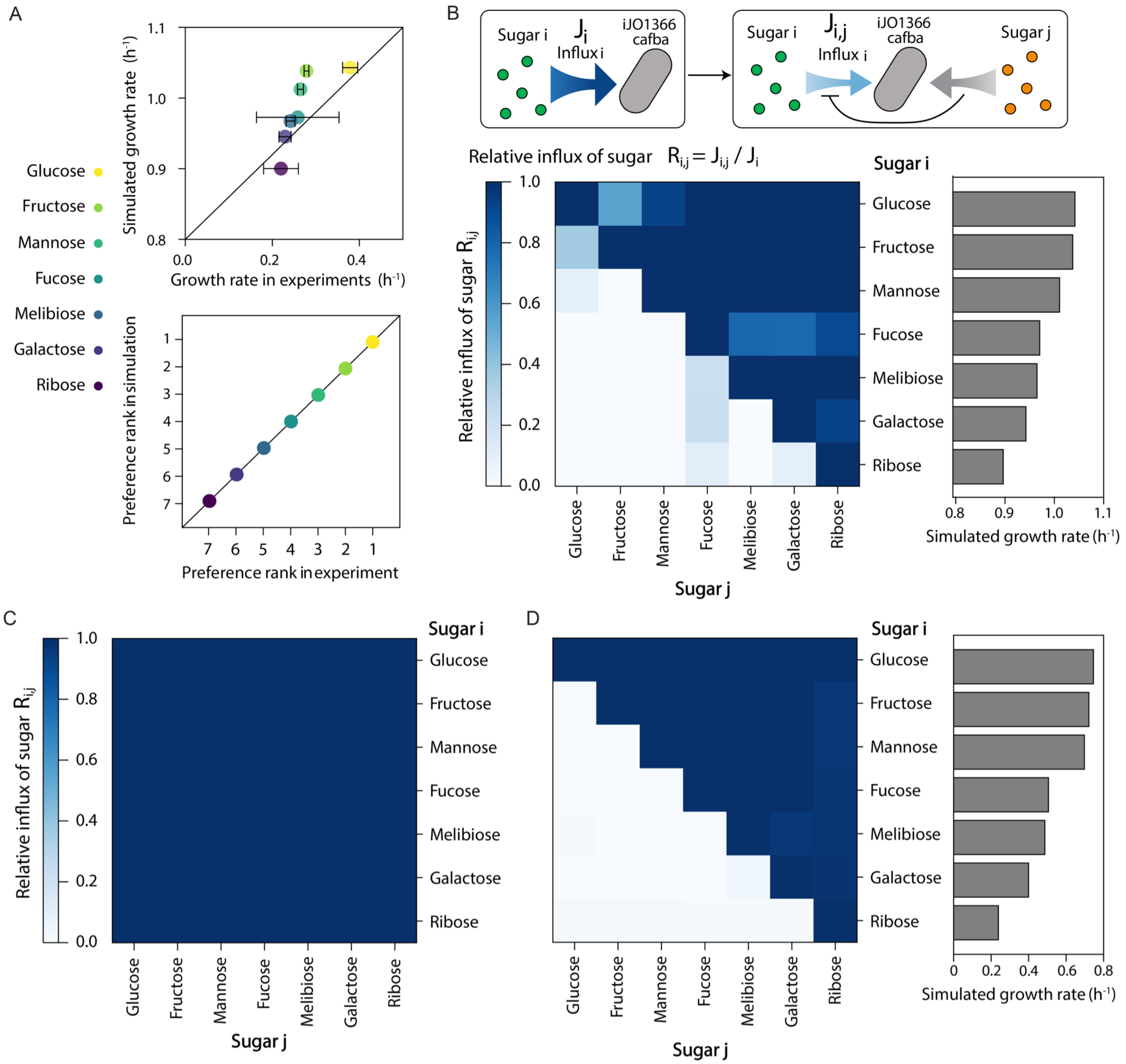


**Fig. S1 A metabolic hierarchy in sugar utilization order is consistent with the hierarchy in growth rate.** (A) A scatter plot of growth rate on 7 sugars and their ranks in experiments or the simulation. Error bars in the horizontal axis indicate standard deviations of the empirical growth rate (Methods). (B) The metabolic influx of 7 sugars was simulated using the model with the global resource allocation constraint (CAFBA, Methods) in the absence or presence of each of the other sugars. We first computed the influx of sugar *i* in the absence of other sugar ($J_i$) , which results in the maximum influx of sugar *i*. Then, we simulated how much the influx of sugar *i* was reduced when another sugar *j* (orange) is present and available for the model ($R_{i,j}$). If $R_{i,j}$ does not change (≈ 1.0), sugar *i* is preferred to sugar *j*. On the other hand, if $R_{i,j}$ decreases, sugar *i* is less preferred than sugar *j*. We displayed the relative influx of each sugar as a heatmap. We set concentrations of two sugars as equivalent per amount of carbon atoms (setting the lower bound of each exchange reaction as -120 mmol×gr$^{-1}$×hr$^{-1}$×Catoms$^{-1}$). Rows and columns are ordered by the hierarchy of growth rate. (C) Simulated metabolic influx of 7 sugars in iJO1366 model without the global allocation constraint (CAFBA). The relative influx of each sugar in the presence or absence of other sugar is displayed as a heatmap. (D) A typical example of the suppressed metabolic influx of sugars depending on the availability of the other sugars. We simulated the influx of each of 7 sugars using the evolved model (i.e. randomly adding or deleting reactions 10000 times). The heatmap shows the relative influx of each sugar in the presence or absence of other sugars. Sugars were ordered by their growth-rate rank.

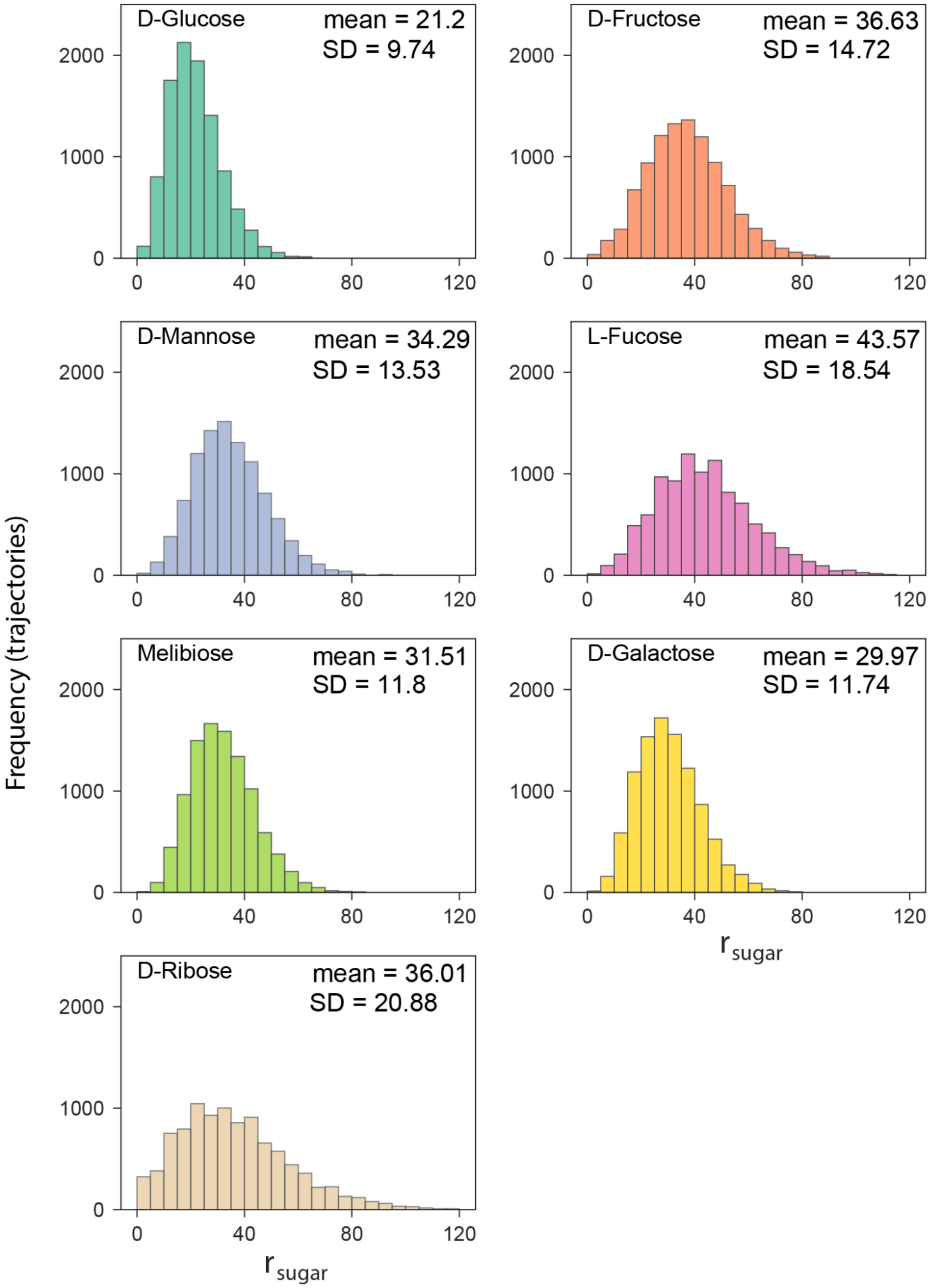

**Fig. S2 Large variation in propensity of rank flips of 7 sugars among evolutionary trajectories.** Total rank flips during random walks in each sugar ($r_{sugar}$) in 10000 evolutionary trajectories is displayed as a histogram. The mean and standard deviation (SD) values are displayed in each panel.

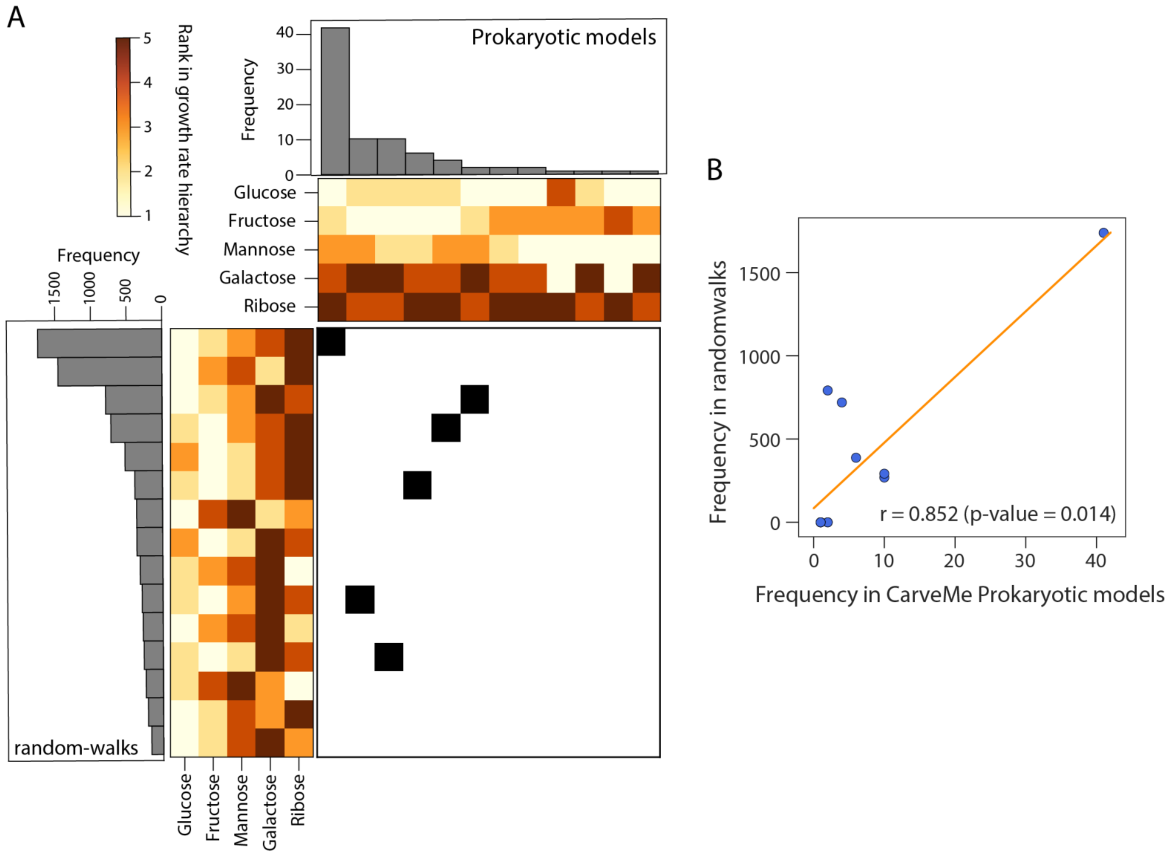


**Fig. S3 Similarity in sugar preference ranks after random walks to those observed in bacterial metabolic models.** We analyzed the rank in growth-rate of sugars using metabolic models constructed by CarveMe. We selected metabolic models that can use at least 5 of the sugars as a sole carbon source. This resulted in a total of 81 metabolic models belonging to 3 phyla, 17 families, and 40 genera. We display their preference ranks (panel A, top heatmap) in order of frequency (histogram). We display a similar diagram for the top 15 most frequent hierarchies in random walks (N = 9974) in the left hand side panel. We found 6 of the 15 most common hierarchies in the random walks, representing ~40 % (4201/9993) of the trajectories, corresponding to the 6 most frequent hierarchies in real organisms' models. (B) Correlation in the frequency of rank configurations between real organisms' models and random walks. For all the rank configurations observed in the CarveMe models (top panel in A), we plotted the frequency in the 81 prokaryotic models and 10000 random walks (N = 9974). Orange line is a linear regression curve, *r* stands for the Pearson's correlation between those two parameters. P-values is obtained using a permutation test (Methods).

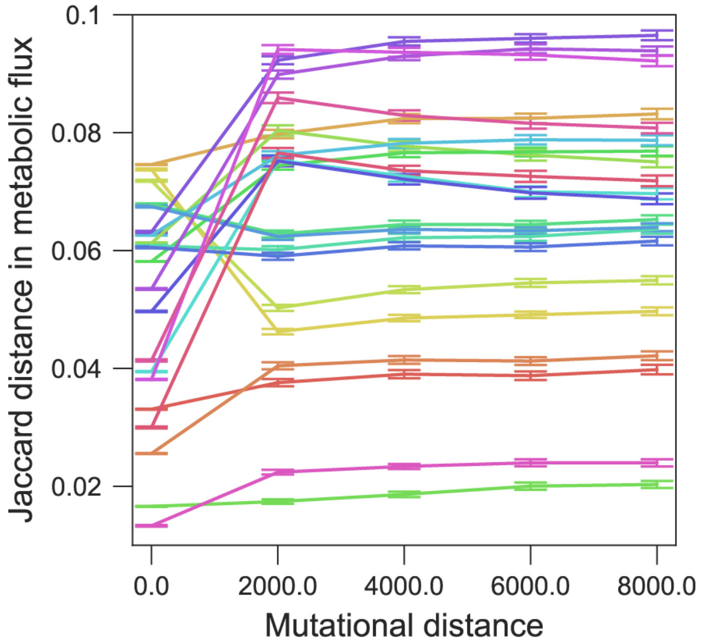


**Fig. S4 Convergence of metabolic dissimilarity between pairs of sugars during the random walks.** Using the same approach as Fig. 2C, we explored temporal changes in metabolic dissimilarity in 21 pairs of sugars during a random walk. We randomly picked up 1000 evolutionary trajectories and computed pairwise Jaccard distance in processing metabolic pathways every 2000 mutations (Methods). Error bars indicate standard error of the mean (SEM).

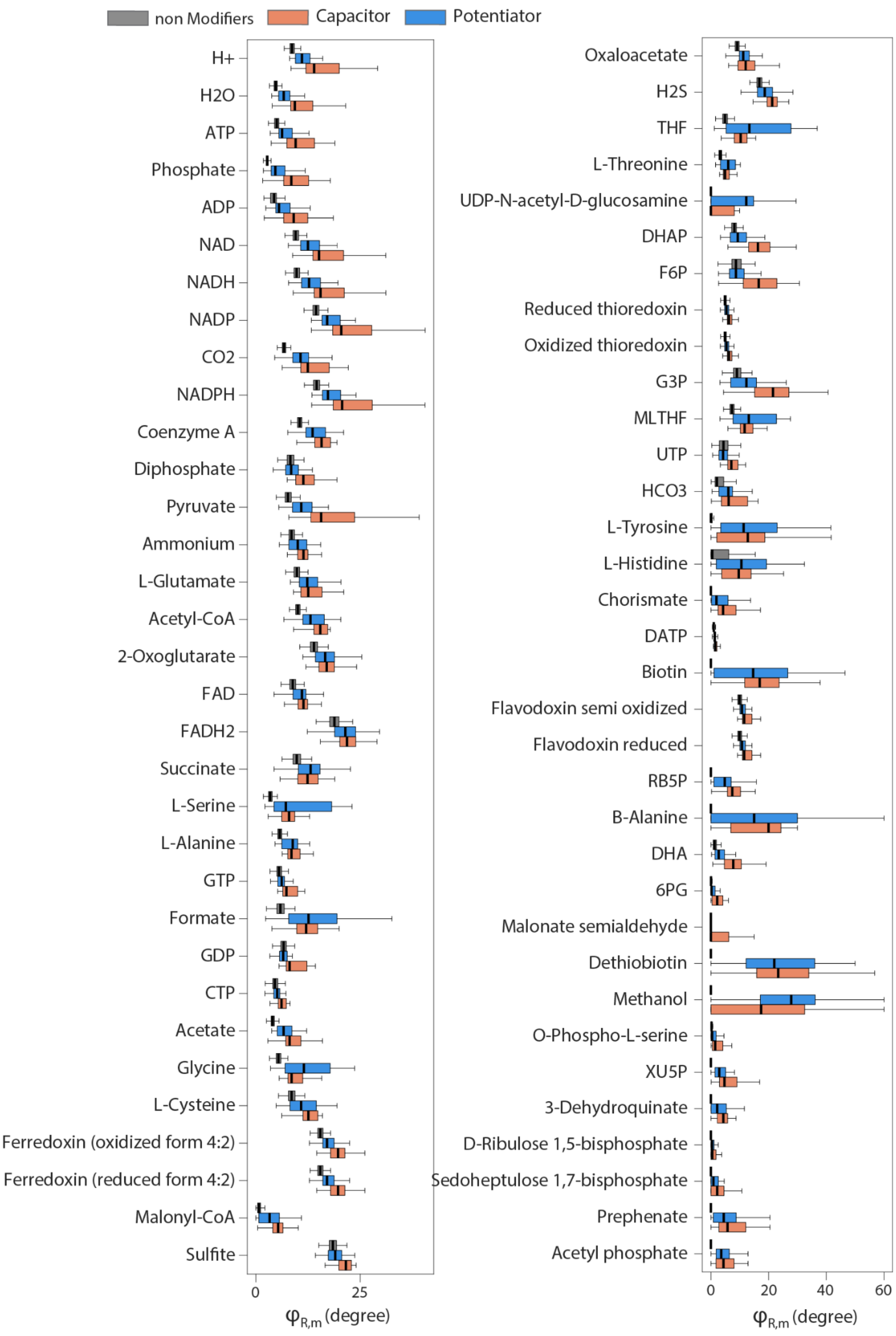


**Fig. S5 Susceptible metabolites by mutations on evolutionary modifiers.** We show metabolites whose flux is significantly more susceptible to perturbations in capacitors or potentiators, compared to non-modifiers (p-value < $10^{-6}$, Wilcoxon rank-sum test with FDR correction).

Metabolites were sorted by the connectivity (i.e., the number of other metabolites connected by a single reaction). The average flux sensitivity levels against the mutations of each reaction group ($\varphi_{R,m}$, Methods) are shown as box plots .

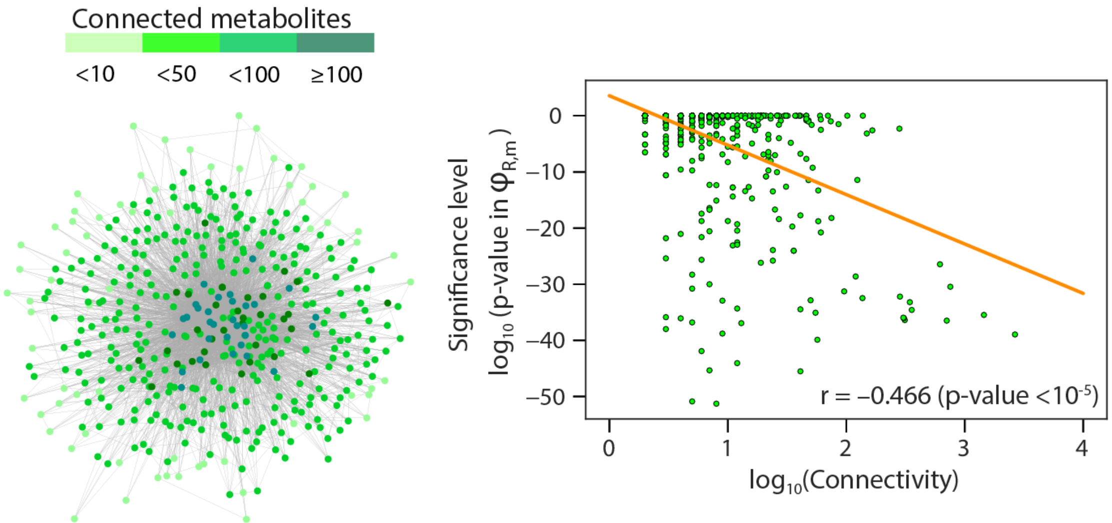


**Fig. S6. Flux sensitivity to modifiers correlates to the connectivity in the network.** We checked how the connectivity of metabolites (i.e.. the number of other metabolites in the network connected by a single reaction) affects the difference in $\varphi_{R,m}$ between modifiers and non-modifiers. In the left panel, we show a graph of the network in a bacterial universal metabolic model. Each dot stands for a metabolite, and a pair of connected metabolites (i.e., involved in common reactions) are linked by a gray line. The connectivity is represented by color. In the right panel, we plotted the number of connected metabolites against the statistical significance level of $\varphi_{R,m}$ in the modifiers compared to the non-modifiers. Pearson's correlation is displayed with p-value in the permutation test (n = 100000).

### Supplementary Text

#### Sugar metabolic hierarchy emerges from the constrained allocation model

A basic framework of metabolic simulation in this study consists of simple stoichiometric and thermodynamic constraints such as mass-balance and consistency with thermodynamic reversibility, and optimal flux distributions are usually determined by linear programming to maximize biomass production rate by FBA only taking into account those constraints. Given that sugar preference rank (i.e. sequential consumption of sugars one after another) is suggested to be a product of optimal protein resource allocation on the mixed carbon sources (Mori et al.), incorporation of the scheme of resource partitioning into the basic constraint-based genome-scale metabolic models scheme is necessary for reproducing this phenomenon. A recent study presented a new FBA framework, called Constrained allocation flux balance analysis (CAFBA), which incorporates costs and optimal allocation of proteomes to biological processes such as ribosome-affiliated, transport, and biosynthesis by single additional global constraints.

Despite the simplicity of the implementation, CAFBA allows us to quantitatively reproduce cellular growth strategies (e.g. over-flow metabolism as a byproduct of optimal proteome allocation).

We first tested whether the preference for sugars emerges in metabolic models using the CAFBA scheme. Here, we tested a simple case: the *E. coli* model can utilize two types of sugar in the environment, and the concentration of those two sugars is equivalent. Then we studied how much uptake (influx) of one sugar is suppressed by the other competing sugar using *iJO1366* incorporating CAFBA constraint (called *iJO1366*-CAFBA hereafter, see Materials and Methods). We focused on the 7 sugars, whose rank in growth-rate is consistent with those observed in the experiment. We analyzed mixtures of 7 sugars (21 pairs of them) in *iJO1366*-CAFBA (Fig. S1A). In Fig. S1B, the sugars are arranged in order of growth-rate. Weak suppressive effects of uptake by the competing sugars (sugar $j$) are observed on the upper-right triangle in the matrix (i.e. when sugar $i$ is high-ranked than sugar $j$), while strong suppressive effects of uptake are observed on the lower-left triangular region in the matrix (i.e. when sugar $i$ is low-ranked than sugar $j$). Therefore, there are perfect matches in a metabolic hierarchy in the utilization of sugars and their ranks in growth-rate: the uptake of a non-preferred sugar (i.e. supporting lower growth rate) is suppressed in the presence of a preferred sugar (i.e. supporting higher-growth rate). For instance, in the case of glucose, the highest sugar in the rank (supporting the highest growth rate) of all, the presence of other sugars does not affect its uptake rate in most cases (i.e. the influx of glucose does not change compared to that in the absence of other sugars). On the other hand, the uptake of ribose, the lowest sugar in the rank (supporting the lowest growth rate) of all, is strongly repressed to zero. We confirmed that the perfect consistency in the preference rank in the utilization of sugars and their rank in growth-rate can still be maintained after rewiring the metabolic network (Fig. S1C). Thus, we use the rank in growth-rate as a proxy for the preference rank of sugar utilization. Of note, this metabolic hierarchy does not emerge when basic *iJO1366* is used for this analysis (Fig. S1D). In all pairs of the sugars, there is no suppression of uptake rate in the presence of two sugars, indicating that the *iJO1366* model always co-utilizes all of the sugars in the environment, and the sequential utilization requires the implementation of a constrained resource allocation scheme.
